# Dynamic Current Clamp Experiments Define the Functional Roles of *I*_*K1*_ and *I*_*to,f*_ in Human Induced Pluripotent Stem Cell Derived Cardiomyocytes

**DOI:** 10.1101/135400

**Authors:** Scott B. Marrus, Steven Springer, Eric Johnson, Rita J. Martinez, Edward J. Dranoff, Rebecca Mellor, Kathryn Yamada

## Abstract

The transient outward potassium current (*I_to_*) plays a key, albeit incompletely defined, role in cardiomyocyte physiology and pathophysiology. In light of the technical challenges of studying adult human cardiomyocytes, this study examines the use of induced pluripotent stem cell-derived cardiomyocytes (iPSC-CMs) as a system which potentially preserves the native cellular milieu of human cardiomyocytes. ISPC-CMs express a robust *I_to_* with slow recovery kinetics and fail to express the rapidly recovering *I_to,f_* which is implicated in human disease. Overexpression of the accessory subunit KChIP2 (which is not expressed in iPSC-CMs) resulted in restoration of a rapid component of recovery. To define the functional role of *I_to_*, dynamic current clamp was used to introduce computationally modeled currents into iPSC-CMs while recording action potentials. However, iPSC-CMs exhibit action potentials with multiple immature physiological properties, including slow upstroke velocity, heterogeneous action potential waveforms, and the absence of a phase 1 notch, thus potentially limiting the utility of these cells as a model of adult cardiomyocytes. Importantly, the introduction of modeled inwardly rectified current (*I*_*K1*_) ameliorated these immature properties by restoring a hyperpolarized resting membrane potential. In this context of normalized action potential morphologies, dynamic current clamp experiments introducing *I_to,f_* demonstrated that there is significant cell-to-cell heterogeneity and that the functional effect of *I_to,f_* is highly sensitive to the action potential plateau voltage in each cell.

## Introduction

Normal cardiac function requires organized, unidirectional propagation of electrical activity, a process dependent on the activation and inactivation of multiple ion channels resulting in the sequential generation of depolarizing and repolarizing currents.[1] Abnormalities of a key repolarizing cardiac potassium current, the fast transient outward current (*I_to,f_*), encoded by members of the Kv4 family, are associated with both heart failure and Brugada syndrome, a congenital arrhythmia syndrome characterized by precordial ST-segment elevation and a risk of malignant arrhythmias.[2, 3] However, the functional role of *I_to,f_* in normal human cardiac rhythm as well as arrhythmogenesis remains incompletely defined.

*I_to,f_* has several inter-related roles in cardiac myocyte function; it contributes to the action potential (AP) waveform and rate-dependent action potential properties as well as to excitation-contraction coupling by influencing calcium influx.[4–8] The role of *I_to,f_* in shaping cardiomyocyte action potentials varies across species, reflecting species-specific differences in other ion channel properties. In rodents with brief action potentials, *I_to,f_* contributes importantly to rapid repolarization.[1] In large mammals with prolonged action potentials, the role of *I_to,f_* is more complex. The rapid activation and inactivation of the current appears to confine its role to the phase 1 notch of the action potential. However, canine experimental and modeling studies indicate that changes in *I_to,f_* alter *I_Ca,L_*, and thereby exert a complex influence on the AP waveform; this relationship results in a non-linear relationship between *I_to,f_* density and action potential duration.[5, 7, 9]

Due to the complex role of *I_to,f_* in shaping action potential waveforms, it is important to examine the functional role of the current in the context of the other native currents present in cardiomyocytes. The recent development of induced pluripotent stem cell technology and the refinement of methods to drive cardiomyocyte differentiation from pluripotent cells offer a novel and promising avenue for the study of cardiomyocyte cellular physiology. However, the utility of induced pluripotent stem cell derived cardiomyocytes (iPSC-CMs) and embryonic stem cell derived cardiomyocytes (ESC-CMs) naturally hinges on how closely they recapitulate adult human cardiomyocyte physiology.

Broadly speaking, pluripotent stem cell derived cardiomyocytes (PSC-CMs) appear to most closely resemble embryonic or immature cardiomyocytes. Over the course of *in vitro* differentiation, iPSC-CMs and ESC-CMs develop many features of mature cardiomyocytes, including expression of cardiac specific transcription factors, contractile proteins, and excitation-contraction coupling.[10] However, the morphology of the cells remains immature, sarcomeres are disorganized, and t-tubules are absent.[11] Furthermore, iPSC-CMs exhibit action potentials with a wide variety of waveforms, generally interpreted as a mixture of ventricular, atrial, and nodal action potential waveforms.[12] However, the action potential exhibit several properties which are not typical of mature cardiomyocytes, including spontaneous diastolic depolarization, slow upstroke velocity, and prolonged action potential duration.[11]

Previous studies have examined the ionic conductances which underlie the action potential waveforms.[12] To date, thorough characterization of the biophysical properties of all of the ion channels expressed in iPSC-CM remains incomplete, but the properties of the ion channels appear to be largely similar to the ion channels expressed in adult ventricular myocytes albeit with a few notable exceptions, including a smaller magnitude inwardly rectifying current (*I*_*K*1_) and a larger magnitude funny current (*I_f_*).[11]

In this study, we demonstrate that iPSC-CMs fail to exhibit *I_to,f_* due to the absence of the essential subunit KChIP2 whereas the closely related but kinetically distinct current *I_to,s_* is robustly expressed.

Although spontaneous action potentials in iPSC-CMs exhibited highly heterogeneous and immature features, iPSC-CM action potential morphology was rendered essentially mature with the introduction of a computationally modeled *I*_*K*1_. This maneuver provided a reasonable model of adult human cardiomyocytes action potentials in which the functional role of *I_to,f_* could be examined. Using dynamic current clamp, we demonstrate that the effect of *I_to,f_* on action potential waveforms varies from cell to cell and is markedly sensitive to the context of other ionic conductances in the cell.

## Materials and Methods

### IPSC generation and cardiomyocyte differentiation

Induced pluripotent stem cells were generated as previously described.[13] Briefly, primary fibroblast lines were established from both a commercially available foreskin fibroblast line (ATCC) (line BJFF) and dermal fibroblasts isolated from a skin biopsy of a healthy 36 yo male (line F10336). Each line was transduced utilizing the non-integrating CytoTune-iPS Sendai Reprogramming Kit (A1378001, Life Technologies). Individual iPSC lines were generated by manual picking and expansion of individual clones in a feeder-free defined system (mTeSR1, StemCell Technologies) and Matrigel (354277, Corning). A minimum of two clones were frozen down for each line and characterization was performed for karyotype, pluripotency and tri-lineage germ layer analysis. All lines had a normal karyotype and expressed endodermal, mesodermal, and epidermal transcripts after spontaneous embryoid body differentiation (S1–S3 Fig). One clone each from neonatal fibroblasts and adult dermal fibroblasts were used for further studies. Written consent was obtained from all participants; the process for consent and acquisition of adult dermal fibroblasts was approved by the Human Research Protection Office at Washington University School of Medicine (protocol #201104172).

Cardiomyocyte differentiation was carried out as previously described using small molecule inhibitors of GSK3 and Wnt signaling; optimal concentrations for both inhibitors were determined based on percentage of beating cells.[14] IPS cells were passaged using accutase (Innovative Cell Technologies). One well of a 6-well plate was seeded with 1x10^6^ iPS cells in mTeSR1 with 5 μM ROCK inhibitor (Y27632, Tocris) and grown to confluence. On Day 0 of the differentiation, cells were fed 4 ml per well with RPMI media supplemented with B27 without insulin (Gibco/Life Technologies). GSK3 inhibitor (CHIR99021, Tocris, 12 μM for BJFF.6 and 6 μM for F10336.6) was added to the media on Day 0. On Day 1, precisely 24 hours after addition of CHIR99021, cells were fed with RPMI+B27 without insulin. On Day 3, 2 ml of conditioned media per well were mixed with 2 ml fresh RPMI+B27 without insulin; the Wnt signaling inhibitor IWP2 (Tocris, 5 μM for BJFF.6 and 10 μM for F10336.6) was added. On Day 5, cells were fed with RPMI+B27 without insulin. On Day 7, and subsequently every 2–3 days, cells were fed with RPMI+B27 with insulin.

On day 20 of differentiation, non-cardiomyocytes were eliminated by feeding with DMEM without glucose (Gibco) supplemented with 4 mM lactate for 3–5 days.[15] Between day 30 and 50 of differentiation, cells were harvested for analysis. Cardiomyocytes were digested with two 10 minute incubations with 0.02% EDTA (Sigma) followed by a brief (~3 min) incubation in 0.25% trypsin; digestion was halted with the addition of RPMI+B27 supplemented with 10% fetal bovine serum. After low-speed centrifugation, cells were replated at low density on plates prepared with 0.1% gelatin in RPMI+B27 with 10% FBS. Cells were fed with RPMI+B27 every 2–3 days and used for electrophysiological recording at least 5–7 days after digestion.

## Electrophysiological recording

To confirm cells’ identity as cardiomyocytes, the mitochondrial dye TMRM (5-10 nM, Molecular Probes, Life Technologies) was added to each plate 15 minutes prior to recording and only cells exhibiting red fluorescence were studied.[16] For experiments involving overexpression of td-tomato, cells to be recorded were identified on the basis of morphology. Recording pipettes fashioned from borosilicate glass (WPI, TW150F-4) had resistances of 2–3 MΩ when filled with recording solution. Whole-cell K+ currents were recorded at room temperature (20–23°C) using previously described solutions and protocols.[17, 18] Action potentials were recorded at 37°C using pipettes containing (in mM): K aspartate 76; KCl 20; MgCl_2_ 2.5; HEPES 10; NaCl 4; CaCl_2_ 6; EGTA 10; K_2_ATP 10; and NaGTP 0.1. Calculated free calcium was 73 nM and free magnesium was 0.36 mM (calculated with maxchelator.stanford.edu). Experiments were controlled by and data were collected using an Axopatch 1D, digitized with a Digidata 1322A, and stored on a PC running Clampex 9.2 (Molecular Devices).

## KChIP2 overexpression

The construction and generation of adenoviral vectors expressing the human KChIP2 subunit and td-tomato under the control of an inducible ecdysone promotor have been previously reported. ([19, 20]; a gift from Jeanne Nerbonne, Washington University School of Medicine). Briefly, a bi-cistronic adenoviral shuttle vector expressing td-tomato and human KChIP2 separated by an internal ribosomal entry site in a single transcript under the control of the ecdysone promoter was used to generate adenovirus. For overexpression experiments, ~3e9 viral particles were added to 2 ml culture media in a 35 mm plate; in addition, ~3e8 viral particles of a second virus (AdVgRXR) encoding the ecdysone receptor which binds muristerone A were also added to the culture media. Two hours after infection, the culture media was supplemented with 0.5mM muristerone A. After ~24 hours, cell expressing td-tomato were identified for electrophysiological recordings. Control experiments were performed using a virus expressing only td-tomato.

## Dynamic current clamp

Using the same recording conditions as above for action potentials, the cell membrane voltage was simultaneously digitized with a Digidata 1322A for storage by Clampex 9.2 and digitized with a NI PCI-6010 DAQ card (National Instruments) installed on a separate PC running a 64-bit real-time Linux kernel (RTAI). A BNC-2110 (National Instruments) connected the amplifier output to the input of the DAQ card. Simulated currents were computed based on the cell membrane voltage (sampled at 10 kHz) using RTXI software[21] and the output of the simulation converted to an analog signal by the PCI-6010 DAQ card. The RTXI output and the Clampex output (used to deliver a brief, depolarizing action potential trigger at 1 Hz) were combined using a custom-made voltage summing junction and the combined signal delivered to the external command of the amplifier. The liquid junctional potential was calculated using Clampex 9.2 and added to the measured transmembrane voltage signal prior to performing the modeled current calculations. Equations for the computational models (see supporting data) of human ventricular *I*_*K*1_ and *I_to,f_* were previously published by O’Hara et al.[22]

## Transcript analysis

Total RNA was isolated using the Qiagen RNeasy Mini Kit (74104) with DNase treatment. Reverse transcription and SYBR Green real time PCR were carried out as previously described.[23] Primers used for RT-PCR are summarized in Table 1. RNA expression levels were normalized first to GAPDH and second to troponin (to normalize for the variable cardiomyocyte content of each differentiation). Tissue from non-failing human hearts was obtained from donor hearts deemed unsuitable for transplantation. Immediately after explantation, small pieces of tissue were placed in RNAlater (Life Technologies) for at least 24 hours, after which RNAlater was removed and the tissue frozen at –80°C until use. Tissue was obtained from donor hearts deemed unsuitable for transplantation; written consent was obtained from the next-of-kin according to the policies of Mid-America Transplant Services. The study was approved by the Human Research Protection Office at Washington University School of Medicine (protocol #201107099).

**Table 1.**
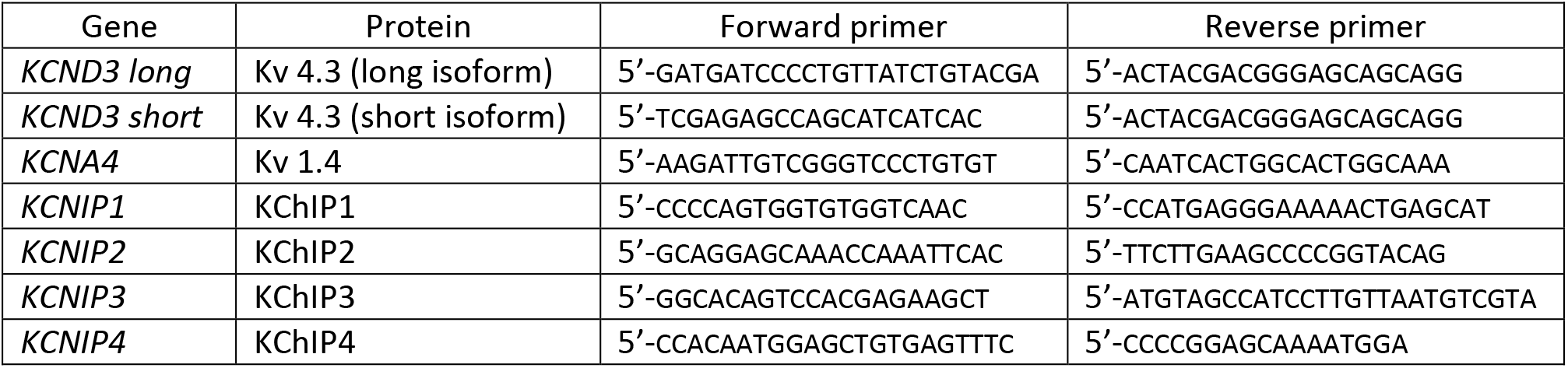
Sequence-specific primers used in SYBR Green RT-PCR

## Data analysis

Analysis of electrophysiological data was performed using MatLab (Mathworks). Whole-cell membrane capacitance was calculated by integrating the capacitative transient elicited by ±5 mV steps from a −70 mV holding potential. The decay phase of the elicited current was fit with a single exponential function; in cells with a biphasic decay, a two exponential function was used and the relative contribution of each component calculated from the weighting of each exponent, as previous described[24]. Action potentials were analyzed for resting membrane potential, peak upstroke velocity, the presence and magnitude of a phase 1 notch, and action potential duration. All values are reported as mean±SEM. Statistical analyses were performed using Prism (GraphPad). Comparisons between groups were performed using either unpaired t-tests or 1-way ANOVAs with Dunnett’s multiple comparisons test.

## Results

### IPSC-CMs express a transient outward potassium current

All IPSC-CMs (n=39) exhibited a rapidly activating, rapidly inactivating transient outward potassium current (Fig 1 and S4 Fig). The transient outward current decay exhibited a fast component with a time constant (τ_inact_) of 74.9±5.4 ms (at 40 mV), similar to values reported for adult LV cardiomyocytes (~45–65 ms).[25, 26] A subset of cells (20 out of 39) also exhibited a more slowly inactivating component (τ_inact_ 635±163 ms) which has not been described in adult human LV cardiomyocytes. Subsequent analysis therefore focuses on only the fast-inactivating component of the transient outward current. The current density of the rapidly inactivating current, *I_to_*, was 6±0.9 pA/pF (at 40 mV), a value similar to the magnitude of the transient outward current reported in adult LV cardiomyocytes (~2–10 pA/pF).[25, 26]

**Fig 1.**
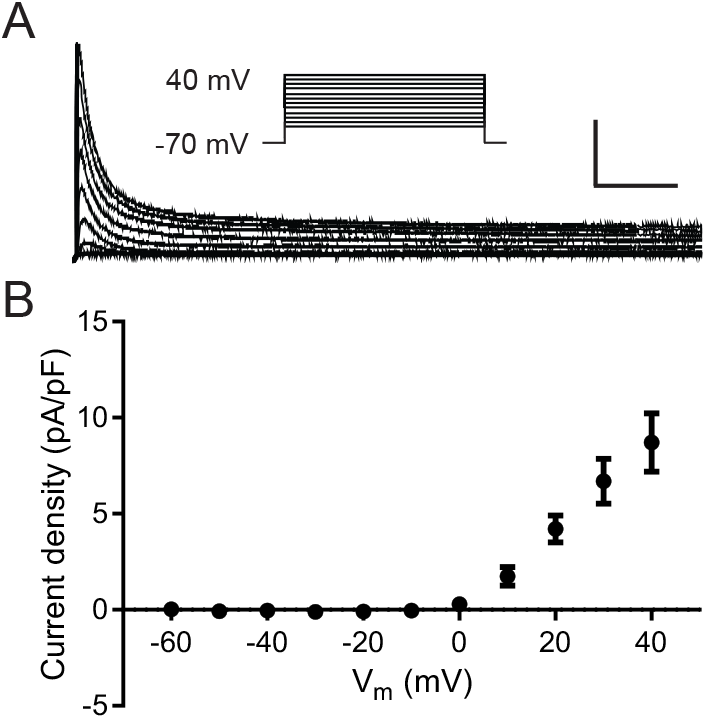
iPSC-CMs (line BJFF) express a rapidly activating and inactivating transient outward current. (A) Representative whole-cell Kv currents (normalized to cell capacitance) recorded from iPSC-CMs in response to voltage steps from 40 mV to −10 mV in −5 mV increments from a holding potential of −70 mV. Capacitative transients have been removed for clarity. Scale bars: 5 pA/pF, 200 ms. (B) Current-voltage relationship for fast-inactivating component (calculated as peak current minus steady state current) of the whole cell current recorded in iPSC-CMs (n=39, mean±SEM).

### *I_to_* in ISPC-CMs exhibits slow recovery kinetics

Two phenotypically (and molecularly) distinct transient outward currents have been described in various species, a fast *I_to_* (*I_to,f_*) characterized by rapid recovery from inactivation (on the order of 10s of ms) and a slow *I_to_* (*I_to,s_*) characterized by slow recovery from inactivation (on the order of 1000s ms).[24, 27] *I_to_* in iPSC-CMs exhibited predominantly slow recovery consistent with (*I_to,s_*) (Fig 2 and S5 Fig), with a time constant of 3735±352 ms. In a minority of cells (2/14), a faster component of recovery (although slower than reported for human *I_to,f_*) was also evident with a time constant of 298±213 ms.

**Fig 2.**
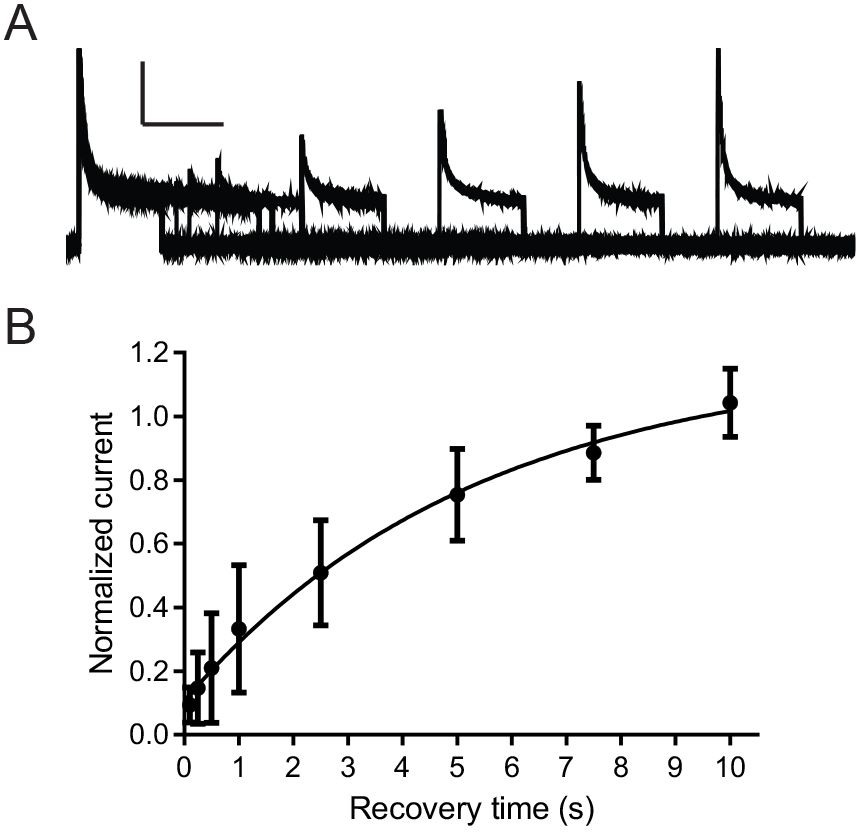
*I_to_* in iPSC-CMs (line BJFF) recovers slowly from steady state inactivation. After inactivating the current during 1.5 sec. prepulse steps to +40mV, cells were hyperpolarized to −70 mV for times ranging from 0.1 to 10 sec. before a second depolarization to +40 mV was applied to assess the extent of recovery. (A) Typical whole-cell Kv current recorded during the inactivation recovery protocol. Scale bars: 5 pA/pF, 1 sec. (B) The amplitude of *I_to_* after each recovery interval was normalized to the amplitude of the current during the prepulse. The time course of inactivation recovery (mean±SEM at each timepoint) is best fit with a single exponential function with a time constant of 3735±352 ms.

### Expression of KChIP2 subunit restores *I_to_* rapid recovery kinetics

Transcripts for the pore-forming subunits *KCND3* (long and short splice variants) encoding Kv4.3 and the KCNA4 transcript encoding Kv1.4, underlying *I_to,f_* and *I_to,s_*, respectively, are expressed at comparable levels in both iPSC-CMs and adult left ventricular tissue (Fig 3 and S6 Fig). However, the KCNIP2 transcript encoding the accessory subunit KChIP2 was expressed at significantly lower levels in iPSC-CMs compared to adult left ventricular tissue (Fig 3 and S6 Fig). KChIP2 has been previously shown to be an essential component of *I_to,f_* in murine hearts.[23] Thus, the low expression level of KChIP2 transcript suggests a reason for the lack of rapidly recovering *I_to_* whereas Kv1.4 likely underlies the observed, slowly recovering *I_to_*.

**Fig 3.**
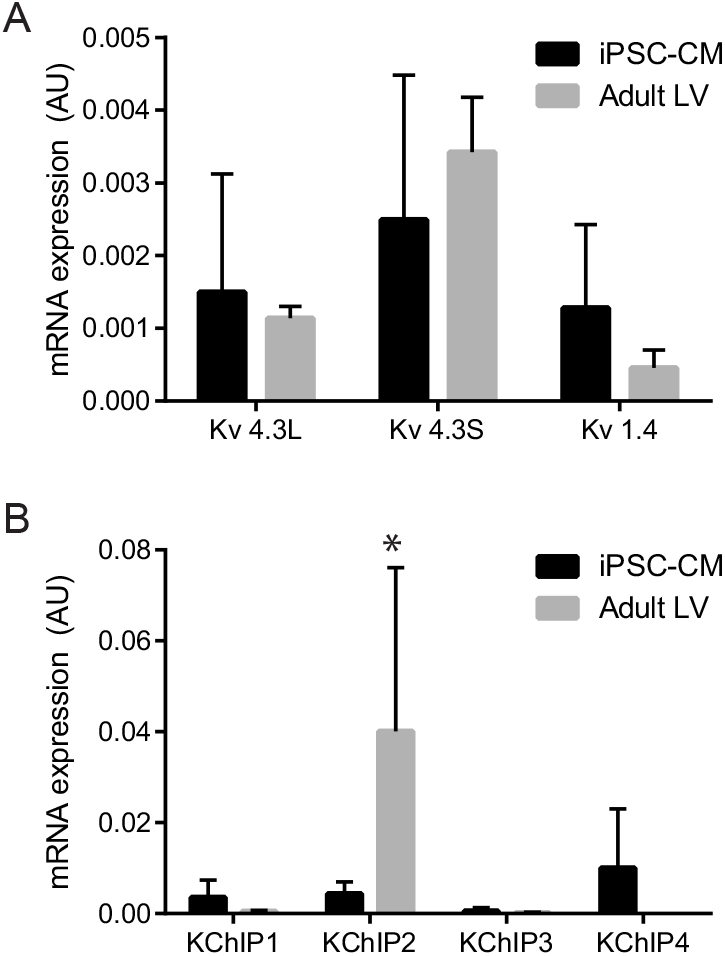
Expression of transcripts encoding subunits underlying *I_to_* in human iPSC-CMs (line BJFF). Transcript expression levels were measured in individual samples of iPSC-CMs (n=6–10) and adult LV apical tissue (n=3), normalized to GAPDH to control for sample loading and subsequently normalized to troponin-T to control for variable numbers of cardiomyocytes in iPSC-CM samples. (A) No significant differences exist between iPSC-CMs and adult human ventricular tissue in transcript expression levels of the pore-forming subunits Kv 4.3L (long splice variant), Kv 4.3S (short splice variant), or Kv1.4 (B) KChIP2 is the predominant KChIP subunit expressed in adult LV tissue and is expressed at significantly (*p=0.005) lower levels in iPSC-CMs; expression in both cell types of transcripts for KChIP1,2, and 4 were negligible.

To directly test the contribution of KChIP2 to *I_to,f_* in human iPSC-CMs, KChIP2 was overexpressed in iPSC-CMs. Overexpression of KChIP2 resulted in an increase in *I_to_* density from 5.9±0.9 pA/pF to 14.1±2.4 pA/pF (p=0.0003). In the presence of KChIP2, 40±5% of *I_to_* recovered rapidly (τ=98±8 ms) while the remaining current recovered slowly (τ=3713±224 ms, p=NS compared to recovery in the absence of KChIP2) (Fig 4 and S7 Fig). The time course of rapid recovery is similar to the value previously reported for *I_to,f_*.[24–26]

**Fig 4.**
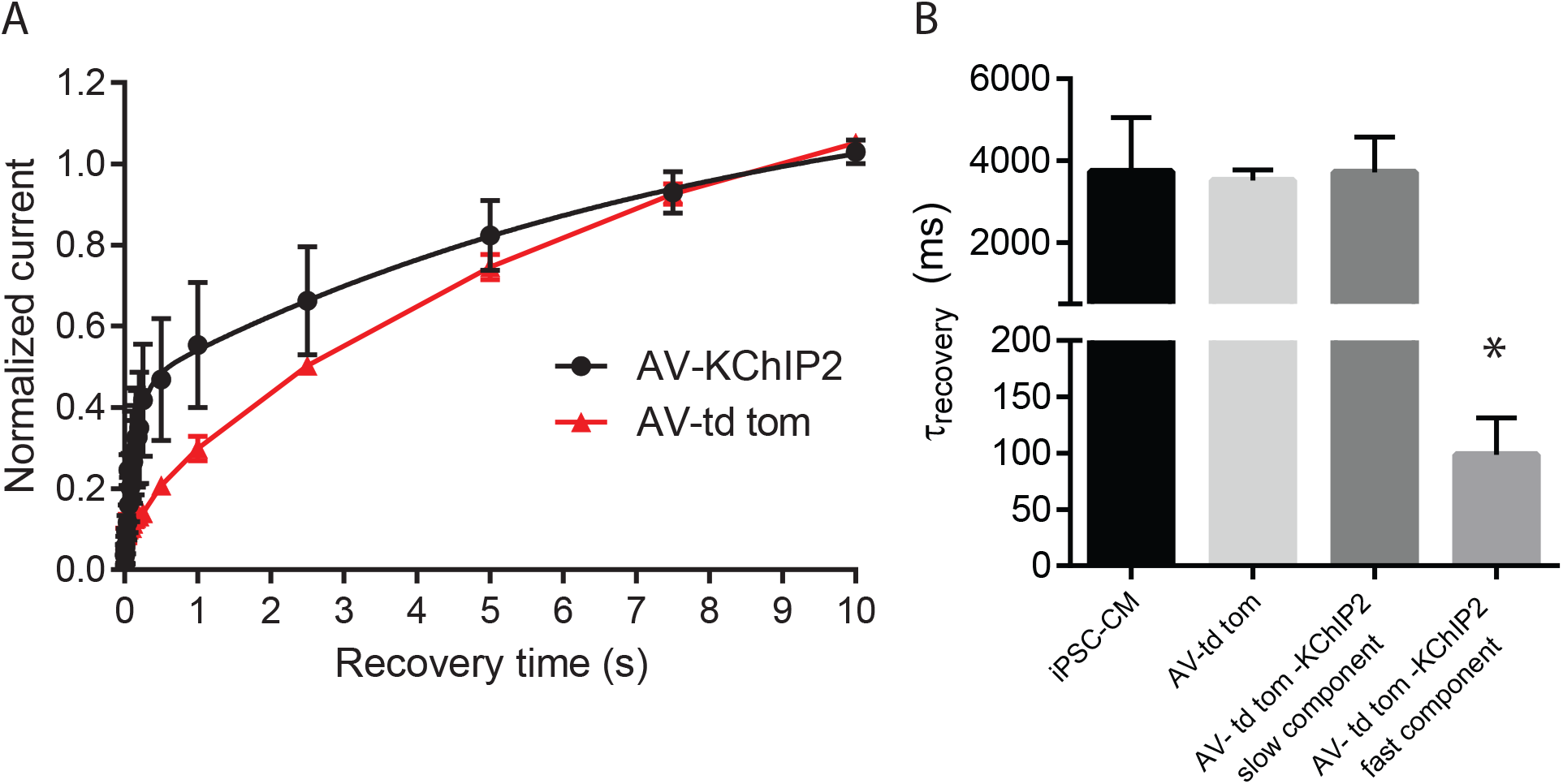
Overexpression of KChIP2 in iPSC-CMs (line BJFF) results in a rapid component of *I_to_* recovery from inactivation. (A) The amplitude of *I_to_* after each recovery interval was normalized to the amplitude of the current during the prepulse with overexpression of KChIP2 and td-tomato (AV-KChIP2) or overexpression of td-tomato only (AV-td tomato). The timecourse of inactivation recovery (mean±SEM at each timepoint) is best fit with a bi-exponential function (compare to Fig 2B) with KChIP2 overexpression but a monoexponential function with overexpression of td-tomato only. (B) After overexpression of KChIP2 or td-tomato alone, the time constant of the slow component of inactivation recovery remained unchanged (first three columns) but overexpression of KChIP2 resulted in an additional, significantly faster component (*p=0.0003) of inactivation recovery (fourth column).

### IPSC-CMs exhibit a wide range of spontaneous AP waveforms

The majority (40/51) of iPSC-CMs exhibited either spontaneous or evoked action potentials. Approximately half (28/51) of the iPSC-CMs recorded exhibited spontaneous action potentials. These cells exhibited a wide range of minimum diastolic potentials (MDP) (−77±2 mV), peak upstroke velocity (dV/dt) (63±13 V/s), and action potential duration (APD90) (627±81 ms) (Fig 5). Spontaneous APs could be grouped into nodal-like, embryonic atrial-like, and embryonic ventricular-like cells on the basis of action potential waveforms.[28] Nodal-like cells (n=4) exhibited an upstroke velocity <10 mV/ms (5±1 mV/ms). Cells with an upstroke velocity > 10 mV/ms were further identified as ventricular-like (n=20) or atrial-like (n=4) on the basis of the presence or absence, respectively, of a plateau phase. The presence of a plateau phase was quantified as the ratio of the slope of the AP waveform between APD70 and APD90 and between APD30 and APD50 (referred to here as the “plateau ratio”); a ratio greater than 1.5 indicated a plateau phase with a more shallow slope than the terminal repolarization.

**Fig 5.**
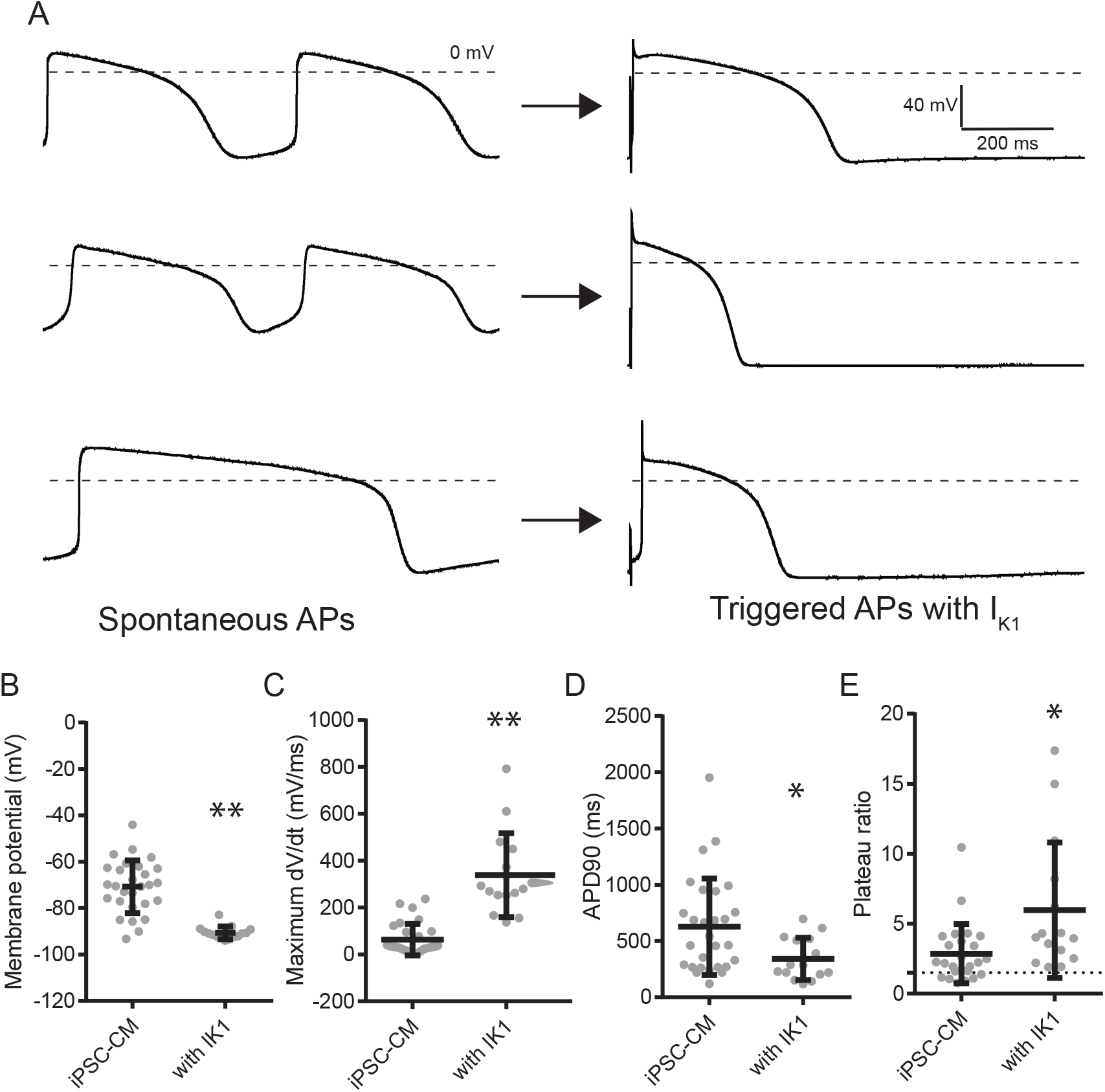
Introduction of computationally modeled *I*_*K*1_ using dynamic current clamp significantly normalizes action potential waveforms (iPSC line BJFF). (A) Representative waveforms of spontaneous action potentials and evoked action potentials after addition of *I*_*K*1_; each pair of traces are from the same cell. The top row demonstrates spontaneous action potentials with a typical atrial waveform; the middle row demonstrates spontaneous action potentials with a typical nodal waveform; the bottom row demonstrates a spontaneous action potential with a ventricular waveform. (B-E) Values from individual cells are represented by grey circles; mean±SEM are indicated by the superimposed bars. Addition of *I*_*K*1_ results in a significantly hyperpolarized resting membrane potential (B), increased maximum upstroke velocity (C) and decreased APD90 (D). E. After addition of *I*_*K*1_ all cells exhibited a plateau ratio > 1.5 (indicated by the dotted line) indicative of a ventricular action potential waveform. **p<0.0001, *p<0.05

### Addition of computationally modeled *I*_*K*1_ results in mature action potential parameters

A prominent feature of iPSC-CMs is a depolarized membrane potential, attributed at least in part to a reduced inwardly rectified current *I*_*K*1_ [11] resulting from reduced expression of the underlying Kir2.1. Dynamic current clamp was used to introduce a computationally modelled *I*_*K*1_ (the mathematical formulation of human cardiac *I*_*K*1_ published by O’Hara et al. [22] was used; S8 Data) at the current density reported in adult human cardiomyocytes which resulted in the cessation of spontaneous action potentials and modification of action potential parameters (Fig 5). The addition of modeled *I*_*K*1_ resulted in significant membrane hyperpolarization compared to the spontaneous MDP (−90±1 mV vs. −69±2 mV, p<0.0001) (Fig 5). Additionally, the peak upstroke velocity (dV/dt) was increased from 57±11 mV/ms to 313±34 mV/ms (p<0.0001), similar to the value reported in adult human cardiomyocytes. The change in action potential morphology was evident by a decrease in the action potential duration (APD90, 619±74 ms vs. 336±37 ms, p=0.003). Importantly, after the addition of *I*_*K*1_ all action potentials exhibited a ventricular-like waveform with a plateau ratio >1.5 (Fig 5). These results indicate that the addition of modelled *I*_*K*1_ render iPSC-CM action potentials more similar to adult ventricular cardiomyocytes.

Given the expression of a robust *I_to_*, it is striking that only 3/28 cells exhibited spontaneous action potentials with rapid phase 1 repolarization and/or a phase 1 notch. One possible reason for this discrepancy is that, due to the depolarized resting membrane potential (RMP), *I_to_* is mostly inactivated. Indeed, restoration of a hyperpolarized resting membrane potential by addition of modeled *I*_*K*1_ resulted in a phase 1 notch in 9/15 cells.

### Addition of computationally modeled *I_to,f_* elucidates the functional role of *I_to,f_*

After the addition of modeled *I*_*K*1_, iPSC-CMs exhibit action potentials similar to adult human LV myocytes, suggesting that these cells represent an appropriate system in which to examine the functional role of *I_to,f_* on action potential duration and waveform. To examine the functional role of *I_to,f_*, computationally modeled *I_to,f_* (S8 Data; based on [22]) was introduced using dynamic current clamp (while simultaneously also delivering modeled *I*_*K*1_). The modeled current was tailored to individual cell size; the model specified a conductance density of 0.02 mS/μF and the cell capacitance was manually entered into the calculations. As expected, the addition of modeled *I_to,f_* resulted in the appearance of a phase 1 notch (4/9 cells without a baseline notch) or a deeper phase 1 notch (5/9 cells with a baseline notch). The voltage of the action potential plateau (measured 25 ms after the peak) was consistently lower after addition of 11±1 pA/pF modeled *I_to,f_* although this failed to reach statistical significance (18±3 mV vs 7±6 mV, p=0.1). There was, however, a clear relationship between the baseline plateau voltage and the magnitude of the effect resulting from *I_to,f_* addition (Figure 6); cells with a more depolarized plateau voltage exhibited little effect of *I_to,f_* addition whereas cells with a less depolarized plateau voltage exhibited a marked decrease in plateau voltage after addition of *I_to,f_*. The effect of *I_to,f_* on action potential duration was similarly heterogeneous; 3 of 9 cells exhibited no effect while the remaining cells exhibited action potential shortening (APD90 = 75±5% of baseline after addition of 11±1 pA/pF modeled *I_to,f_*). These results demonstrate that the functional role of *I_to,f_* is highly dependent on the precise cellular context.

**Fig 6.**
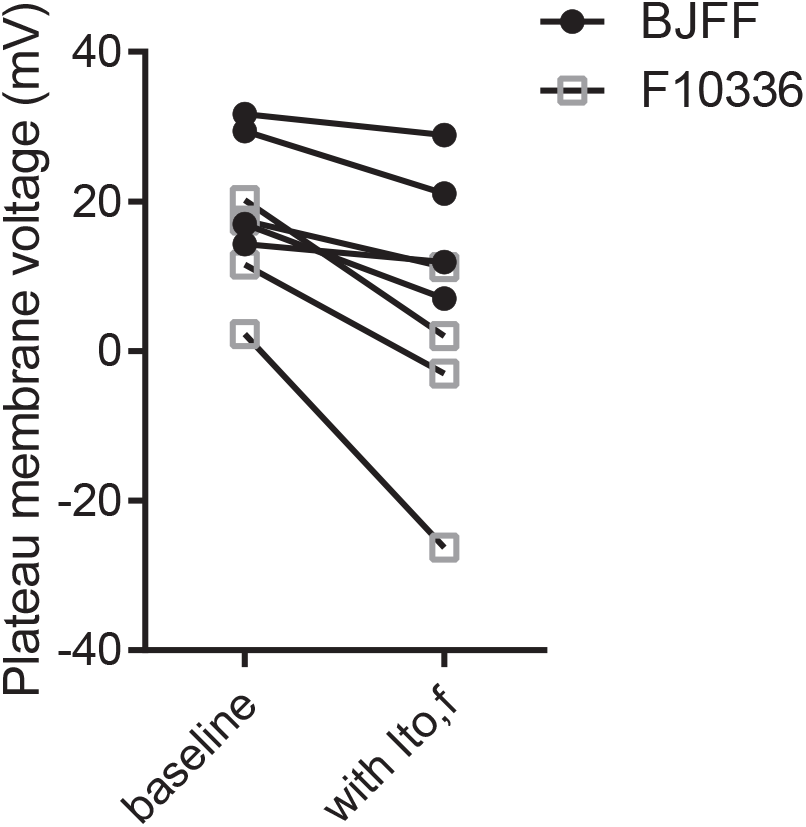
Dynamic current clamp addition of *I_to,f_* to iPSC-CM (lines BJFF and F10336) results in cell-specific changes in plateau membrane potential. (A) Representative action potential waveform (upper panel) at baseline (black) and after the addition of *I_to,f_* (red). The modeled *I_to,f_* is shown in the lower panel. (B) The effect of modeled *I_to,f_* on the plateau membrane potential is greatest in cells with a less depolarized plateau potential at baseline.

## Discussion *I_to_* in iPSC-CMs

The utility of iPSC-CMs as a model of adult ventricular cardiomyocytes naturally depends on the degree to which they recapitulate the electrophysiological properties of adult human cardiomyocytes. Previous work has demonstrated that iPSC-CMs express most of the major ionic conductances present in adult cardiomyocytes and that the time- and voltage-dependent properties of most [12] but not all [29] ionic currents are similar to the currents recorded in adult human ventricular myocytes. However, multiple morphologic and physiologic properties of stem-cell derived cardiomyocytes have been noted to differ from adult cardiomyocytes and are generally interpreted as reflecting a more immature state. [30] Previous characterization of *I_to_* in iSPC-CMs by Cordeiro et al. reported a strikingly slow recovery from inactivation (τ=2380 ms at 37°C, corresponding to approximately 8000 ms at room temperature[25]).[31] In contrast, we observed a recovery timecourse which, albeit slow, was significantly faster than that reported by Cordeiro et al., with a time constant of 3735 ms at room temperature. Of note, Cordeiro et al. used embryoid body differentiation whereas this study used monolayer differentiation; this observation highlights the potential for physiological differences in cells arising from different protocols.

Both this study and Cordeiro et al. found an absence of the accessory subunit KChIP2 in iPSC-CMs; this study further demonstrated that over-expression of KChIP2 resulted in a rapidly recovering component of *I_to_*. It has been previously shown that KChIP2 is an essential subunit underlying *I_to,f_* in the mouse.[23] The present results provide the most direct evidence to date of a similar essential role of KChIP2 in the generation of human *I_to,f_*. Both studies confirm the absence of functional *I_to,f_* in iPSC-CMs, highlighting a limitation of this model for adult cardiomyocytes.

In both canine and rodent hearts, KChIP2 expression increases during post-natal development in parallel with the emergence of *I_to,f_*. [32] Data obtained from neonatal human atrial tissue has also previously demonstrated a post-natal upregulation of KChIP2 corresponds to faster *I_to_* recovery (albeit only by a factor of 2) from inactivation.[33] Taken together, therefore, although it is plausible (and likely) that iPSC-CMs reflect an earlier developmental state, the prominence of an *I_to_* with extremely slow recovery contrasts not only with adult human cardiomyocytes but also with early neonatal human cardiomyocytes.

### Action potential morphology in iPSC-CMs

Multiple previous studies have reported immature action potential properties in iPSC-CMs as well as a depolarized resting membrane potential.[11, 12] Previous theoretical[31] and experimental[34] studies have suggested that restoration of normal resting membrane potential would normalize action potential waveforms. In this study, dynamic current clamp was used to introduce modelled *I*_*K*1_ which restored a normal resting membrane potential, rapid phase 0 upstroke, rapid phase 1 repolarization and reduced the action potential duration. These results complement those of Bett et al.[34] who reported that two distinct AP waveforms (atrial and ventricular) were recorded after introduction of *I*_*K*1_ whereas we observed only ventricular waveforms. One possible explanation for this difference is that Bett et al. studied iCell (Cellular Dynamics) cardiac myocytes (derived from spontaneous differentiation in embryoid bodies) whereas this study used directed cardiomyocyte differentiation. These results highlight the fact that, although introduction of *I*_*K*1_ can largely normalize action potential waveforms, different cardiomyocyte differentiation protocols potentially yield cells with different properties.

### Functional role of I_to,f_

The fast, transient outward potassium current, *I_to,f_*, plays an important albeit complex role in cardiomyocyte physiology and pathophysiology. Computational simulations have predicted that increasing *I_to,f_* conductance approximately 1.5-fold results in moderate action potential prolongation and slightly depolarized membrane potential during the action potential plateau while further increases result in action potential collapse.[7] In contrast, a dynamic current clamp study on canine ventricular cardiomyocytes reported essentially no impact on either plateau voltage or APD; the effect of modelled *I_to_* was confined to the phase 1 repolarization.[9] Here, we show that introducing modeled *I_to,f_* to human iPSC-CMs consistently affects the phase 1 repolarization but that the effect on plateau voltage varies significantly among cells and is sensitive to the baseline plateau voltage (which in turn reflects the baseline conductances in the particular cell). This finding, in conjunction with previously published results, highlight the fact that the functional role of *I_to,f_* is exquisitely sensitive the balance of other ionic currents in the cell and is likely to be cell-type specific.

## Acknowledgements

The authors thank Dr. Michael Pasque for surgical assistance with the collection of human cardiac tissue. The authors thank Translational Cardiovascular Biobank and Repository at Washington University School of Medicine for access to human cardiac tissue.

## Supporting Information

**S1 Fig.**
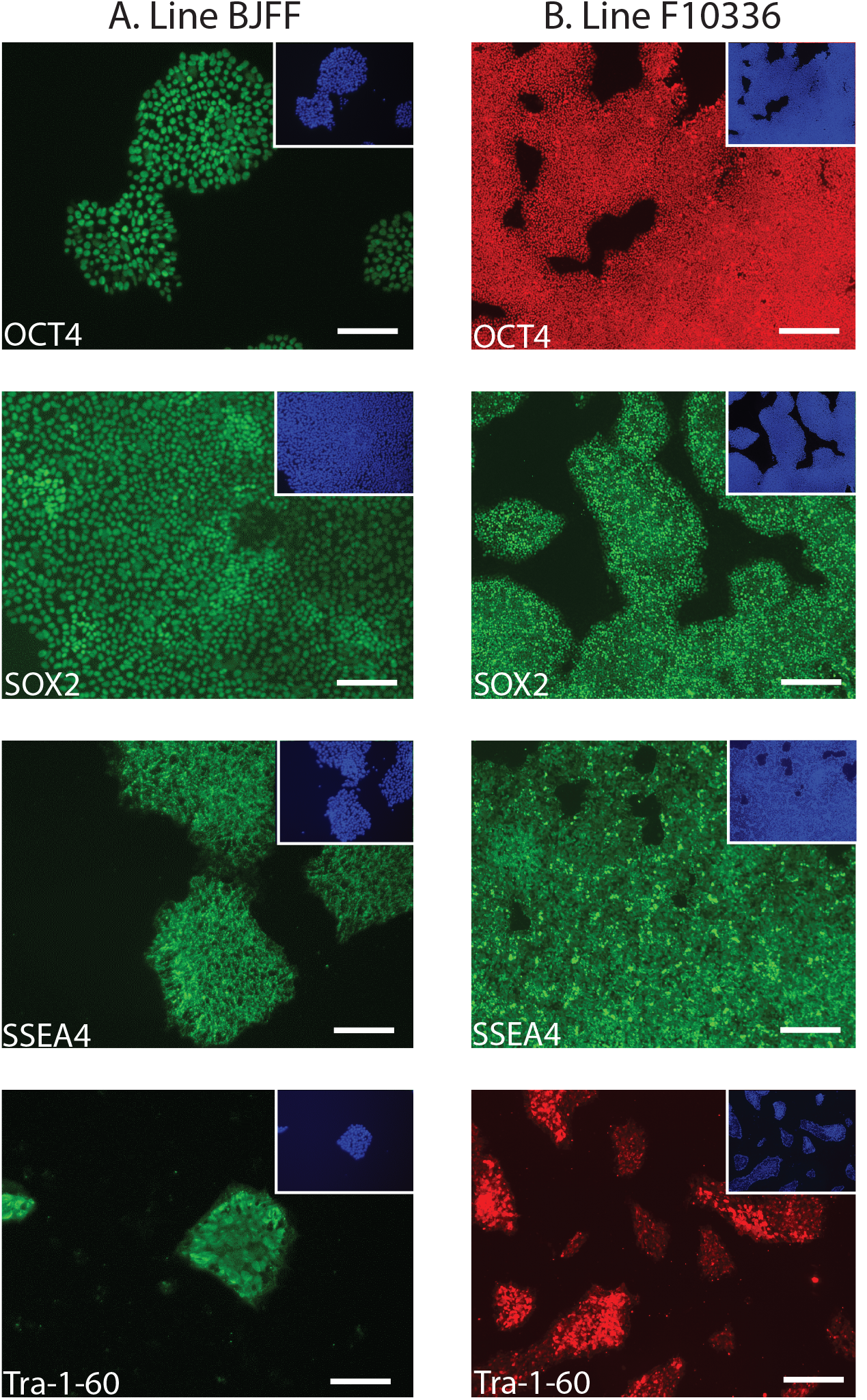
After induction of pluripotency, both iPSC lines express pluripotency markers. IPSCs were plated onto matrigel-coated LabTek 8-chamber slides (Thermo-Fisher 154534PK) and grown in mTesR1 for 48 hours after which they were fixed and stained using The Pluripotent Stem Cell 4-Marker Immunocytochemistry Kit (LifeTechnologies A24881) (line F10336) or the StemLight iPSC Cell Reprogramming Antibody Kit (Cell Signaling Technology, #9092) (line BJFF) according to the manufacturer’s instructions. Immunocytochemistry results demonstrating expression of the pluripotency markers OCT4, SOX2, SSEA4, and Tra-1-60 is shown for line BJFF (column A) and line F10336 (column B). Inset demonstrates nuclear staining using DAPI. Scale bar is 125 μm.

**S2 Fig.**
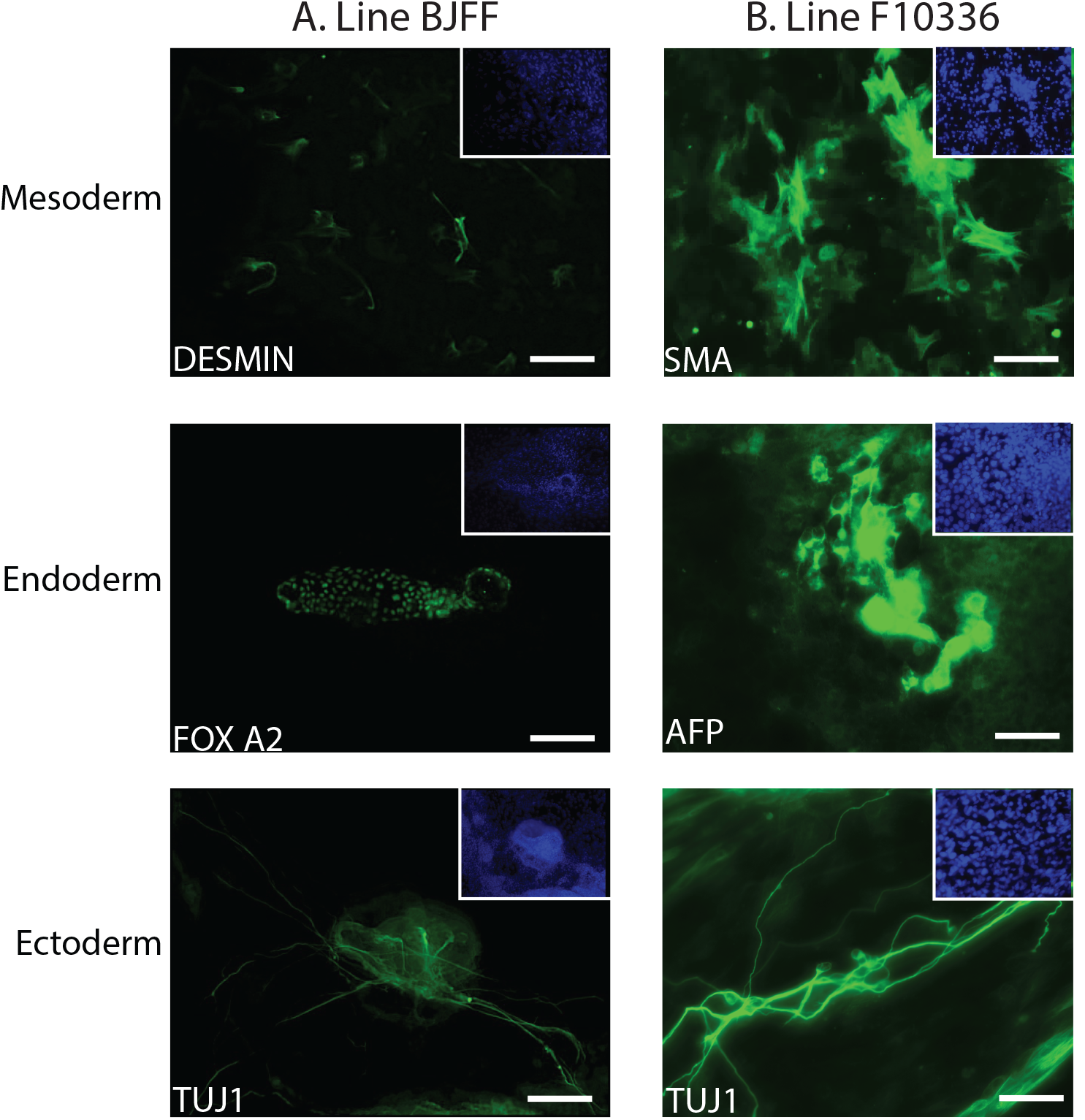
After spontaneous differentiation in embryoid bodies, both iPSC lines express mesodermal, endodermal, and ectodermal markers. EB differentiation was performed by digesting iPSCs with accutase followed by spontaneous EB formation in low attachment plates (Costar 6-well Clear Flat Bottom Ultra-Low Attachment Multiple Well Plates, #3471) for 2–3 days. EB differentiation continued for 3 days after EB formation after which EBs were plated onto 0.1% gelatin coated LabTek 8-chamber slides for an additional 4–6 days. EBs were then fixed and stained using the 3-Germ Layer Immunocytochemistry Kit (LifeTechnologies A25538) according to the manufacturer’s instructions. Immunocytochemistry results demonstrating expression of markers for all three germ cell layers is shown for line BJFF (column A) and line F10336 (column B). Inset demonstrates nuclear staining using Hoecht’s stain. Scale bar is 125 μm.

**S3 Fig.**
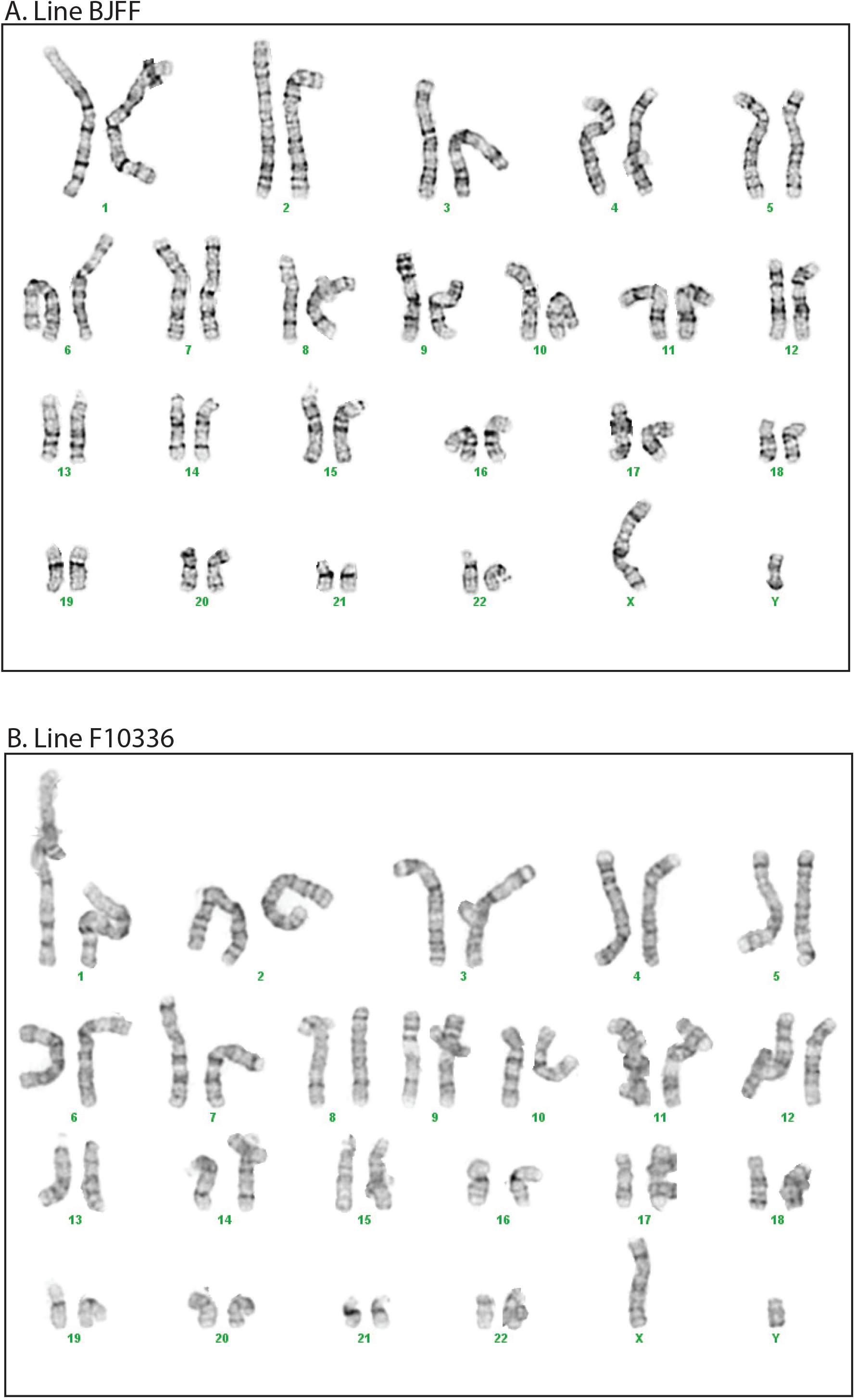
Both iPSC lines have a normal karyotype. Karyotypes were performed by Cell Line Genetics.

**S4 Fig.**
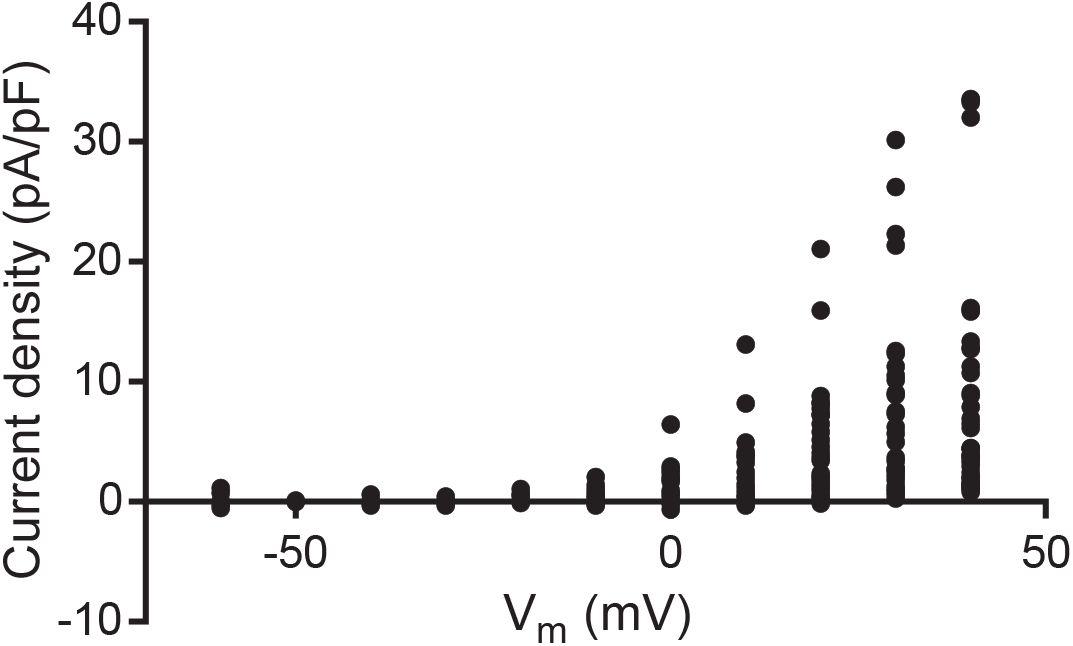
Individual data points for the current-voltage relationship for fast-inactivating component of the whole cell current recorded in iPSC-CMs. For each cell, the magnitude of the fast-inactivating current (calculated as peak current minus steady state current) is normalized for cell capacitance and plotted as a function of voltage.

**S5 Fig.**
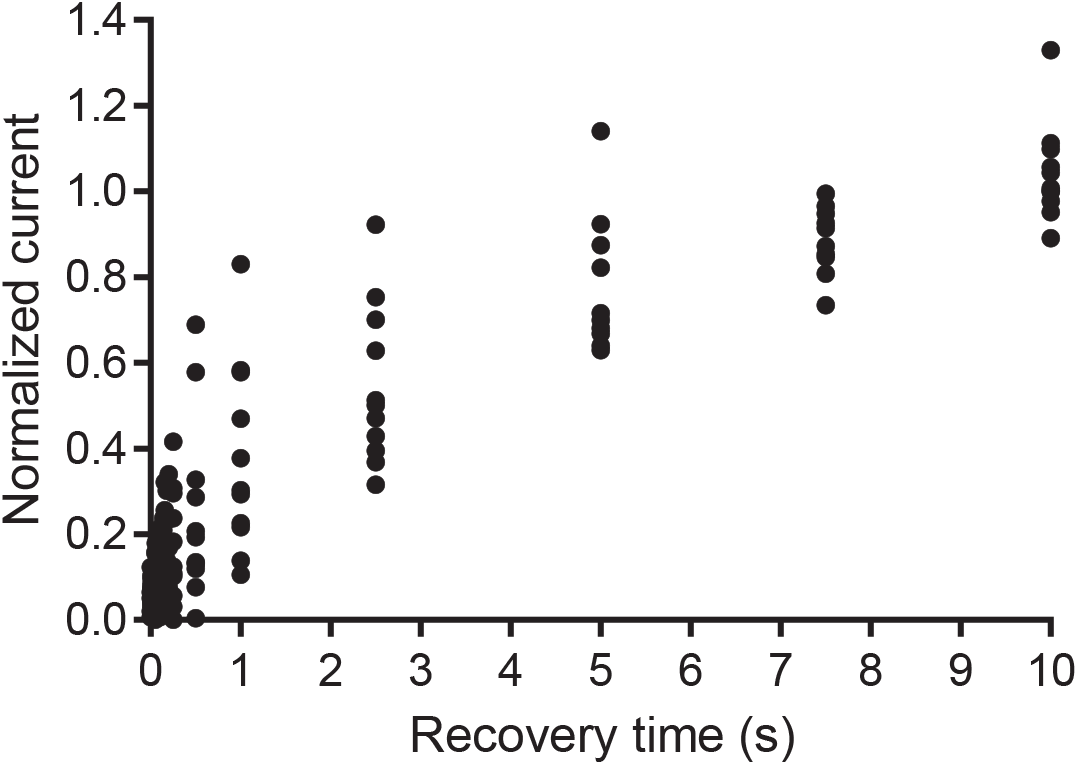
Individual data points for *I_to_* recovery from inactivation. For each cell, the amplitude of *I_to_* after each recovery interval was normalized to the amplitude of the current during the prepulse.

**S6 Fig.**
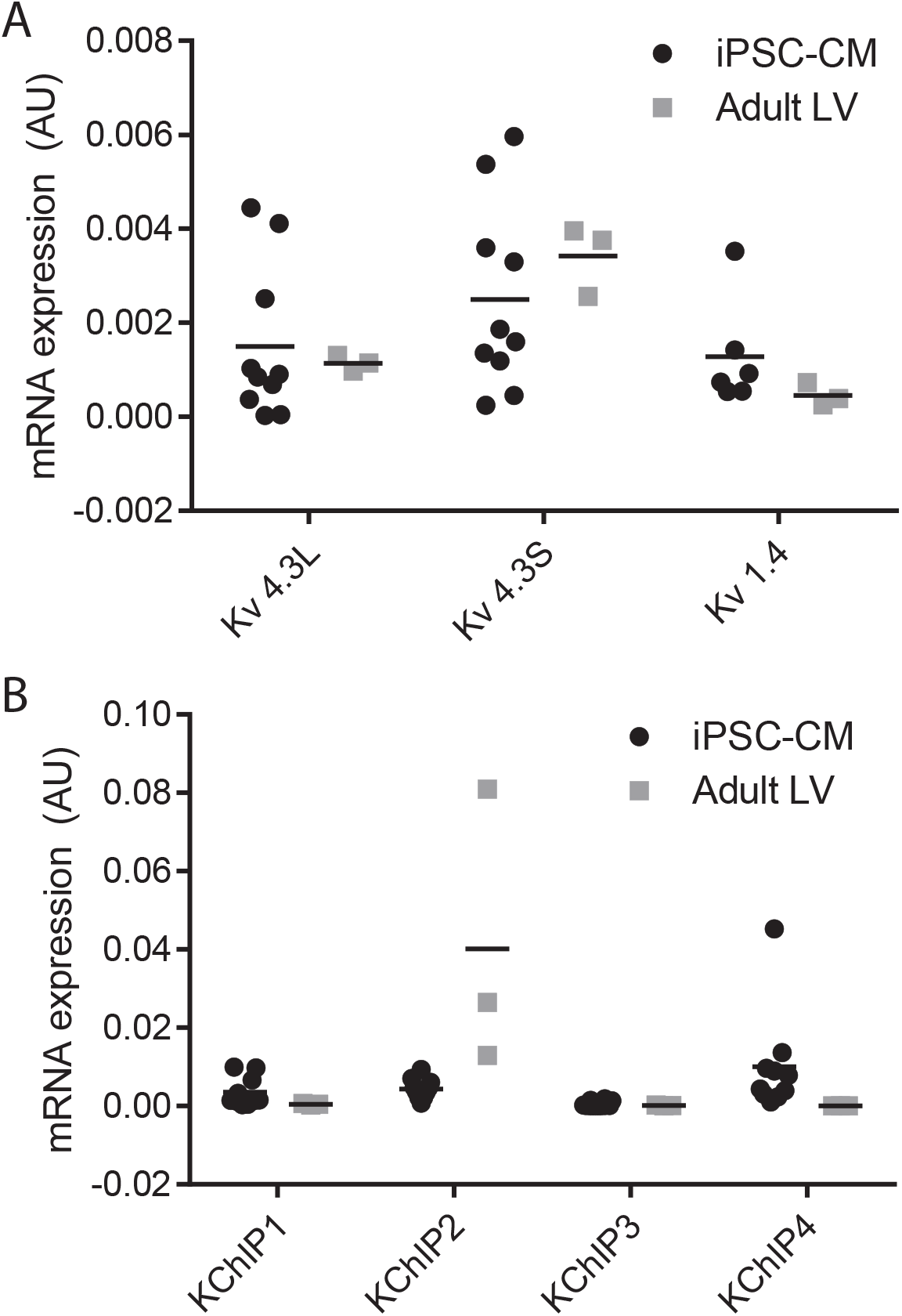
Individual data points for expression of transcripts encoding subunits underlying *I_to_* in human iPSC-CMs.

**S7 Fig.**
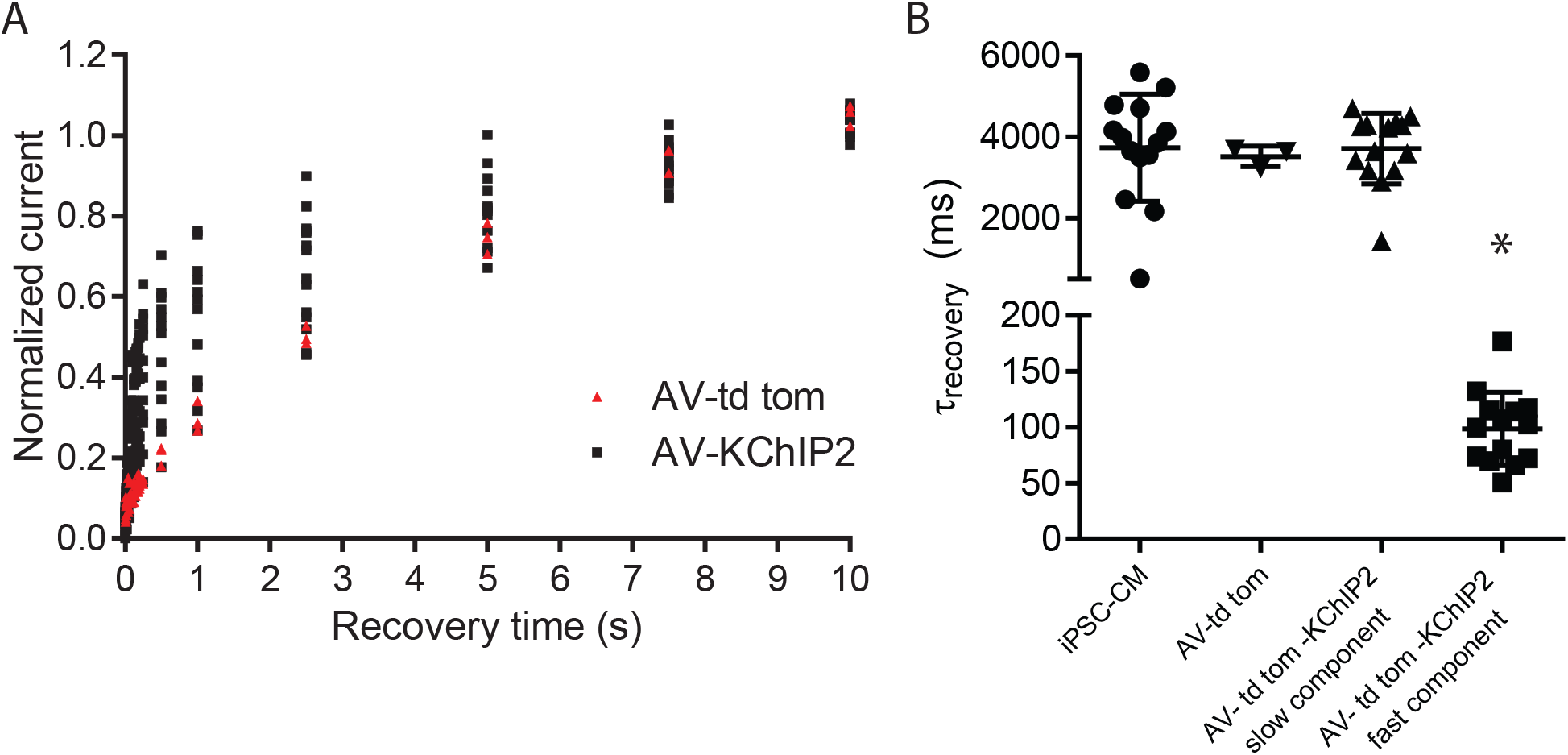
Individual data points for recovery from inactivation with KChIP2 overexpression. (A) For each cell, the amplitude of *I_to_* after each recovery interval was normalized to the amplitude of the current during the prepulse with overexpression of KChIP2 and td-tomato (AV-KChIP2) or overexpression of td-tomato only (AV-td tomato). (B) After overexpression of KChIP2 or td-tomato alone, the time constant of the slow component of inactivation recovery remained unchanged (first three columns) but overexpression of KChIP2 resulted in an additional, significantly faster component (*p=0.0003) of inactivation recovery (fourth column).

**S8 Data**. Formulas for modeled ionic currents in dynamic current clamp experiments.

### Supplementary Data

Computational model for human *I*_*K*1_ used for dynamic clamp experiments using formulas previously published by O’Hara et al.(1) V=membrane voltage, [K^+^]_o_=extracellular potassium concentration, E_k_=reversal potential for potassium, C_m_=membrane capacitance.

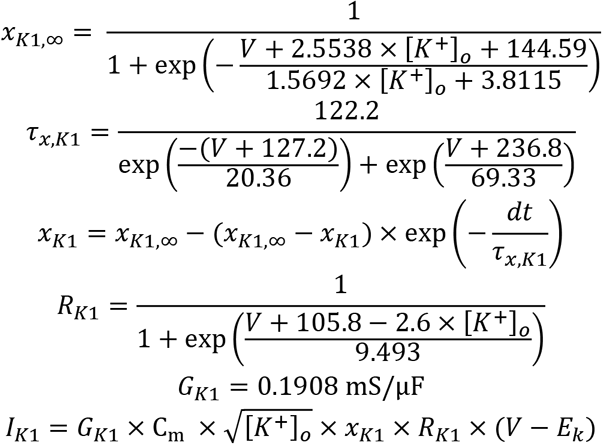

Computational model for human *I_to,f_* used for dynamic clamp experiments using formulas previously published by O’Hara et al.(1)

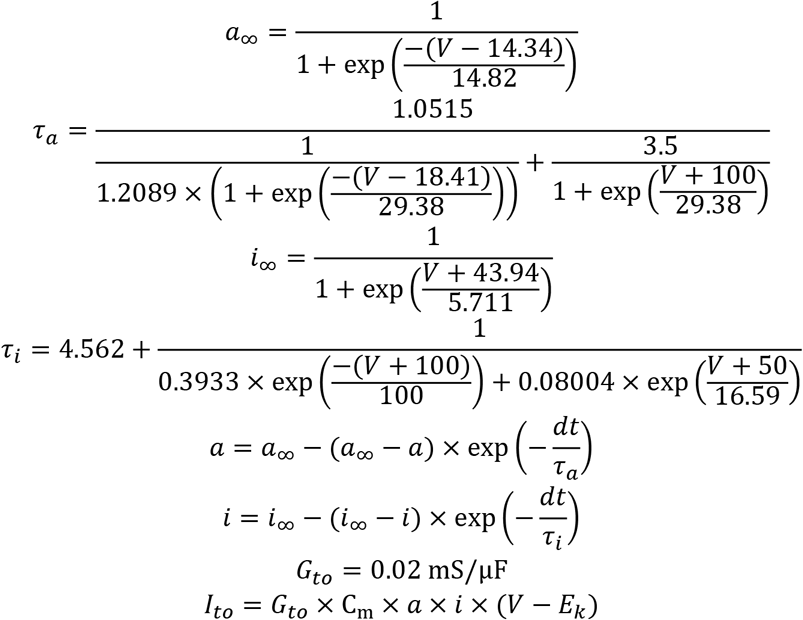

1. O’Hara T, Virag L, Varro A, Rudy Y. Simulation of the undiseased human cardiac ventricular action potential: model formulation and experimental validation. PLoS computational biology. 2011;7(5):e1002061.

